# Descriptors of Pain Quality Associate with Pulpal and Periapical Diagnostic Tests in Patients Experiencing Acute Tooth Pain

**DOI:** 10.1101/262022

**Authors:** O. Erdogan, M. Malek, M. N. Janal, J. L. Gibbs

## Introduction

Dental pain causes a great deal of suffering, stress, and interference in the lives of patients, and relief from pain is a common reason for seeking dental care (Heaivilin et al. 2011). However, how we typically measure pain in dentistry does little to capture the complex experience of pain. Scales such as the visual analog scale (VAS) or numeric rating scale (NRS) are the most commonly used, and allow for the quantification of pain intensity. Used alone, as is most often then case, these scales are quite limited, given the multidimensional nature of pain, with sensory, affective, and temporal dimensions mediated by complex and varied biological mechanisms (Dworkin et al. 2005; von Hehn et al. 2012). Pain qualities (e.g. burning, aching, shooting) have been well studied in relation to clinical diagnosis, and specific descriptors are reported by patients experiencing mechanistically distinct types of pain (Baron et al. 2009). For example, patients with neuropathic pain, the type of pain that occurs after damage or injury to the sensory nervous system (e.g. diabetic neuropathy) will frequently report pain qualities including burning, numb, and/or tingling. Such pain qualities are not often reported in patients experiencing nociceptive or inflammatory pain, which are typically described as aching, throbbing, sharp, or dull. Few studies have evaluated how pain qualities reported by patients experiencing tooth pain relate to the underlying clinical status of the pulp and periapical tissues (Levin et al. 2009).

Although “toothache” is a general term for odontogenic pain, there are multiple mechanisms that can contribute to the clinical presentation. The pulp tissue itself has a remarkably unique innervation, which seems to be hard wired to convey pain with the slightest physical stimulation (Fried et al. 2011). As such, even when the pulp is free of any pathology, pain can occur if the dentinal tubules are exposed and open, as occurs in dentin hypersensitivity. When the dentin is affected by deep caries or fracture, bacteria gain access to the dentinal tubules producing varying degrees of inflammation in the pulp. The severely inflamed or infected vital pulp, is described by clinical entity “symptomatic irreversible pulpitis”, which is characterized by spontaneous pain and mechanical and thermal allodynia and hyperalgesia (e.g., pain on biting, and to cold stimulation of the tooth). If bacteria gain access to the pulp chamber, the pulp tissue progressively becomes necrotic, the sensory afferents that innervate the pulp die back, and the inflammation and infection spread to the periapical tissues. The sensitized nerve endings in the periapical tissues (including periodontal ligament and alveolar bone) will cause mechanical allodynia, where any mechanical stimulation of the tooth painful (Figure 1). Thus diagnostic tests that utilize sensory responses of teeth to thermal and mechanical stimulation are used to infer the biological state of the pulp and periapical tissues, and reach a clinical diagnosis.

**Figure 1.**
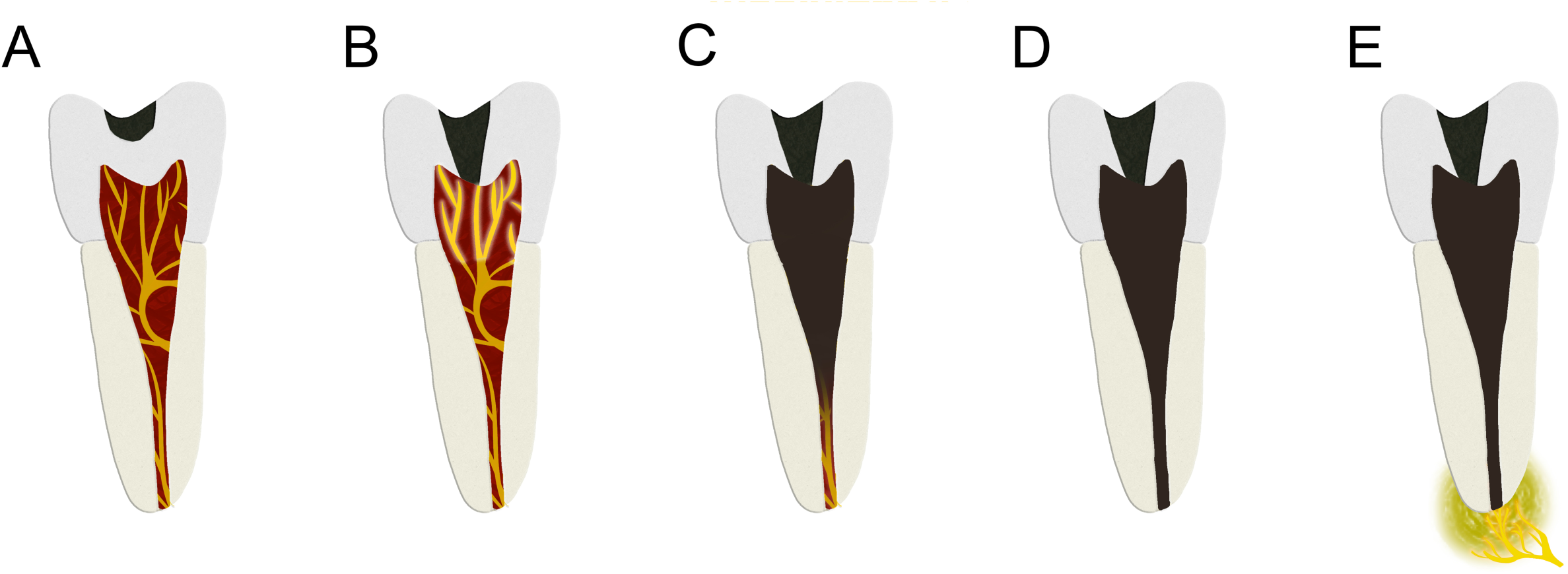
A. The pulp is free of any pathology. Even when the pulp is free of any pathology, pain can occur if the dentinal tubules are exposed and open. B. When the dentin is affected by deep caries or fracture, bacteria gain access to the dentinal tubules producing varying degrees of inflammation in the pulp. The severely inflamed or infected vital pulp is characterized by spontaneous pain and mechanical and thermal allodynia and hyperalgesia. C. Bacteria gain access to the pulp chamber, the pulp tissue progressively becomes necrotic, the sensory afferents that innervate the pulp die back. D. Necrotic pulp. E. The inflammation and infection spread to the periapical tissues. The sensitized nerve endings in the periapical tissues cause mechanical allodynia.

In this study we tested the hypothesis that patients experiencing tooth pain with differing underlying biological mechanisms causing symptoms would report different pain qualities. For example, is pain from a tooth with inflamed and degenerating vital pulp tissues be described differently than pain from a tooth with a necrotic pulp and periapical infection. We tested for an association between mechanical and thermal sensory responses of affected teeth, with pain quality descriptors and pain intensity at rest and in function. Identification of such associations will clarify whether pain descriptors are useful in discriminating the underlying biological processes contributing to dental pain.

## Material And Methods

### Study Overview and Inclusion

This cross-sectional clinical study was approved by the Institutional Review Boards of New York University and University of California San Francisco, which determined the study protocol met the requirements for protection of human subjects. Generally healthy adult men and women who voluntarily presented to the dental clinics for an unscheduled visit due to dental pain were invited to participate in the study, and verbal consent was obtained. To be included in the study patients had to report an intensity of ongoing pain of at least 3 out of 10 on a numerical rating scale. Further, there had to be a clear endodontic or pulpal etiology for the reported pain (e.g. large carious lesion) that was localizable and only involved a single tooth. Patients with generalized severe periodontitis, and non-endodontic odontogenic pain (e.g. pericoronitis) were excluded. Patients with multiple teeth simultaneously causing pain, as well as patients with a history of chronic orofacial pain (including temporomadibular joint disorder and chronic migraine), were excluded from the study. Participation in the study was voluntary, uncompensated, and had no impact on the subsequent dental treatment the patient received.

### Study procedures/Data Collection

Patients completed a verbally administered questionnaire regarding basic demographics, global health, medication usage, pain history and frequency, pain qualities, pain intensity, and pain in function. Specific questions regarding pain qualities were selected based on their prior validation to discriminate between different types of pain (e.g. neuropathic or not) in other instruments including the Standardized Evaluation of Pain (Scholz et al. 2009), The DN4 Scale (Bouhassira et al. 2005), The Neuropathic Pain Scale (Jensen et al. 2005), the LANSS Pain Scale (Bennett 2001), and dental-pain screening questionnaire (Pau et al. 2005). When queried about pain, patients were asked to consider their pain at rest and in function over the past 48 hours. Pain quality questions were formatted as in the following example, “Does your pain feel like a shooting pain?”. If patients responded yes, then a follow up question regarding the intensity of that pain was asked, as in “How intense is your shooting pain?” Patients would respond using a numerical rating scale from 0–10 with 0 described as “no pain”, and 10 described as the “worst possible pain.” The questions were verbally asked to the patient by study investigators and the answers were recorded on an iPad using Google Forms. The resulting dataset did not contain any patient identifiers.

After completion of the questionnaire, standardized endodontic diagnostic testing procedures were conducted by calibrated study personnel, who were either endodontic faculty or residents. A contralateral, adjacent, and the symptomatic tooth were evaluated with each test, in that order. Pulp vitality was first assessed using cold detection. A standard #2 cotton pellet was sprayed with Frigi-dent (Ellman, Hicksville, NY, USA) until saturated. The pellet was placed on the buccal aspect of the tooth, and a new pellet was used for each tooth. Patients reported whether they felt the cold and the intensity of the sensation (none, mild, moderate, severe). Teeth without full coverage restorations were also tested with an electric pulp test. Teeth were dried and the device was set to 6 (SybronEndo Vitality Scanner; Kerr Corporation, Orange, CA, USA). The numerical readout on the device, when the patient experienced sensation was recorded.

For testing of mechanical allodynia of the periapical tissues, the occlusal surface of the tooth was first tapped with mild force using the handle of dental mirror. The unpleasantness of the sensation was recorded was ranked from 1–3 with 1 being mild or no unpleasantness and 2, moderate and 3 severely unpleasant. For palpation testing, the apical region of the root was gently pressed with a gloved fingertip and the sensation was rated using the same 1–3 scale of unpleasantness. Lastly, a bite test was done using a Tooth Slooth (Professional Results Inc., Laguna Niguel, CA, USA). For multicusped teeth, all cusps were tested and the most severe response recorded using the same (1–3) scale. Other clinical parameters recorded included deepest periodontal probing, swelling (yes/no), sinus tract (y/n), and the clinical radiograph was evaluated for the presence of caries (y/n) and periapical radiolucency (y/n). Finally, study personnel who were endodontists evaluated the clinical findings and gave a pulpal and periapical diagnosis for the tooth using the American Association of Endodontists published guidelines.

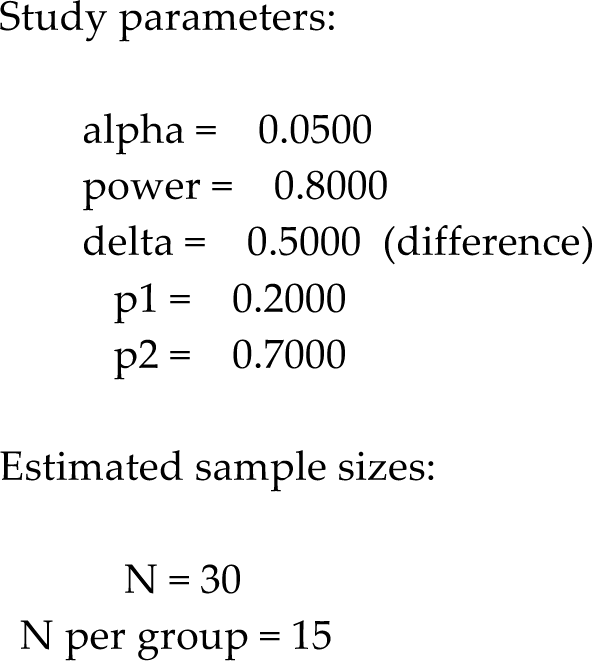

### Statistical Procedures

For this observational study, we used existing studies that measured the frequencies of reporting pain descriptors in patients with different diagnoses, to estimate the needed sample size (Bouhassira et al. 2005). Using the assumption that we would see a 40% difference in the frequency of reporting a descriptor in patients with different diagnoses (e.g. patient with irreversible pulpitis versus pulpal necrosis), and an alpha of 0.05 and power of 0.80, gave an estimation of 46 subjects (23/group) (Calculated using Stata/IC 4.2 for Mac (StataCorp, College Station, TX, USA)). As we planned to test multiple descriptors, we determined that 200–250 patients were reasonable to recruit in the time available for the study and sufficient powered the study.

Data was exported from Google Forms to Excel, and then imported into StataVersion 14 (StataCorp, College Station, TX, USA), which was used to perform organization, labeling, and generation of composite variables. Descriptive analysis of variables was performed to identify frequency, mean and standard deviation. Univariate association between pain descriptor reporting and cold detection and percussion pain test was tested using chi-square analysis. Further, a 2-way ANOVA was carried for the association of selected pain descriptors or global pain measures with independent variables of cold detection and percussion hypersensitivity. A p-value less than 0.05 was considered statistically significant.

## Results

### Patient and Tooth Characteristics

The study sample consisted of 228 subjects who attended a dental emergency clinic due to dental pain. The mean age was 42 (Range of 18–81 (Table 1). A slight majority (53.1%) were female and about half identified as Caucasian (46.9%) (Table 1). Most subjects reported their health to be excellent or good and less than 20% identified themselves as having chronic pain. Most cases had a pulpal diagnosis of either irreversible pulpitis (34.2%) or pulpal necrosis (35.5%) and symptomatic apical periodontitis (67.1%) was the most common periapical diagnosis (Table 1).

**1.** Subject Demographics & Tooth Characteristics

1. Subject Demographics & Tooth Characteristics
| PATIENT-RELATED CHARACTERISTICS |  |
| --- | --- |
| Age, y, mean (SD) | 42 (16) |
| Female sex, n (%) | 121 (53.1) |
| Ethnicity, n (%) |  |
| Hispanic or Latino | 53 (23.2) |
| Non-Hispanic or Latino n (%) | 175 (76.8) |
| Race, n (%) |  |
| White/Caucasian | 107 (46.9) |
| Black/African American | 42 (18.4) |
| Asian | 7 (3.1) |
| Native American/Native Alaskan | 5 (2.2) |
| Hawaiian/ Other Pacific Islander | 3 (1.3) |
| Mixed Ethnic Group | 27 (11.8) |
| None of these | 37 (16.2) |
| General Health n (%) |  |
| Poor & fair | 30 (13.2) |
| Good | 111 (48.7) |
| Very good & excellent | 87 (38.2) |
| Presence of Chronic Pain n (%) | 41 (18) |

| TOOTH-RELATED CHARACTERISTICS |  |
| --- | --- |
| Maxillary Tooth n (%) | 113 (49.5) |
| Tooth Group n (%) |  |
| Incisor | 26 (11.4) |
| Premolar | 52 (22.8) |
| Molar | 150 (65.8) |
| Possible Etiology n (%) |  |
| Caries | 129 (56.6) |
| Fracture | 33 (14.5) |
| Restoration related | 24 (10.5) |
| Reinfection | 14 (6.1) |
| Trauma | 2 (0.9) |
| Other | 21 (9.2) |
| Unknown | 5 (2.2) |
| Pulpal Endodontic Diagnosis n (%) |  |
| Normal Pulp | 5 (2.2) |
| Reversible Pulpitis | 17 (7.5) |
| Irreversible Pulpitis | 78 (34.2) |
| Pulpal Necrosis | 81 (35.5) |
| Previously Initiated Therapy | 19 (8.3) |
| Previously Treated Tooth | 28 (12.3) |
| Apical Endodontic Diagnosis n (%) |  |
| Normal Apical Tissues | 30 (13.2) |
| Symptomatic Apical Periodontitis | 153 (67.1) |
| Asymptomatic Apical Periodontitis | 10 (4.4) |
| Acute Apical Abscess | 31 (14.0) |
| Chronic Apical Abscess | 3 (1.3) |
| Presence of Periapical Radiolucency n (%) | 96 (42.1) |

### Global Pain Measures

On average, the subjects reported a moderate level of pain at the time of evaluation (mean=5.2 on a 10 point numerical rating scale), and severe levels of pain in function (mean= 7.5), and worst pain (mean= 8.5) (Table 2). More than half of subjects had been experiencing this pain for a week or less (58.5%), but note that 21.5% were experiencing pain for more than a month. Although 71.5% of subjects reported using some type of medication for their pain, only 29% had taken medication within the last 4 hours (Table 2). The most commonly taken medications were NSAIDS and opioids. Most patients described the temporal quality of their pain as intermittent (69.8%), meaning they feel pain sometimes but are pain free other times (Jensen et al. 2005).

**2.** Global Pain Intensity

2. Global Pain Intensity
| Pain Intensity* | | Mean Pain Intensity ± SD |
| --- | --- | --- |
| | Pain at present | 5.2±2.9 |
| | Worst pain | 8.6±1.2 |
| | Pain in function | 7.5±2.1 |
| | Pain at rest | 5.2±3.0 |
| Pain Duration |  | N (%) |
|  | 1 week or less | 114 (58.5) |
|  | 1 week up to 1 month | 39 (20) |
|  | 1 month and more | 42 (21.5) |
|  | Missing Data | 33 |
| Temporal Aspect of Pain** |  | N (%) |
|  | Intermittent | 57 (30.2) |
|  | Variable | 89 (47.0) |
|  | Stable | 43 (22.8) |
|  | Missing Data | 39 |
| Pain Medication Use |  | N (%) |
|  | Ever | 163 (71.5) |
|  | Missing Data | None |
|  | Within the last 4 h | 66 (29) |
|  | Missing Data | None |
| Type of Pain Medication Used |  | N (%) |
|  | NSAIDs or other non-opioid combination | 109 (47.8) |
|  | Opioid (including combinations) | 37 (16.23) |
|  | Other drugs such as topical analgesics | 17 (7.46) |
|  | None | 65 (28.5) |

**SD: standard deviation**
**N (%): number of patients (percentage of patients within the population N= 228)**
**\*Pain intensity on 0-10 NRS**
**\*\*Intermittent pain: pain that comes and goes with some periods with no pain; variable pain: constant pain but pain type and intensity changes over time; stable pain: pain that remains constant over time (Jensen et al. 2005).**

### Pain Descriptors

We next determined the frequency and mean pain intensity associated with 22 pain descriptors in subjects experiencing acute dental pain. The pain descriptors used are categorized as paroxysmal/intermittent, continuous, paresthesia/dysthesia, or evoked pain. The most frequently reported descriptors were pain evoked by chewing/biting (88.6%), pain evoked by cold (68%), the continuous descriptors throbbing (77.2%) and aching (78.5%), and the intermittent pain descriptor radiating (67%) (Table 3).

**3.** Pain Descriptors Frequency and Mean & Standard Deviation

3. Pain Descriptors Frequency and Mean & Standard Deviation
| Pain Descriptors | Frequency of Reporting Pain Descriptor (%) | Mean Intensity & Standard Deviation** | Positive Cold Detection n (%) | Painful Percussion n (%) |
| --- | --- | --- | --- | --- |
|  |  |  | 106 (46.5) | 163 (71.5) |
| PAROXYSMAL/<br>INTERMITTENT |  |  |  |  |
| Shooting<br>p | 48.7* | 8.0±1.7 | **62 (55.8)**<br>**0.006** | 81 (72.9)<br>0.63 |
| Stabbing<br>p | 43.9* | 8.1± 1.7 | 48 (48.0)<br>0.69 | 72 (72.0)<br>0.88 |
| Bursting<br>p | 38.6* | 7.8±2.0 | **48 (54.5)**<br>**0.05** | 58 (65.9)<br>0.14 |
| Radiating<br>p | 67.1* | 7.1± 2.2 | 73 (47.7)<br>0.60 | 113 (73.8)<br>0.26 |
| Electric shock-like<br>p | 28.9* | 7.4± 2.0 | 35 (53.0)<br>0.21 | 45 (68.1)<br>0.48 |
| CONTINUOUS |  |  |  |  |
| Throbbing<br>p | 77.2* | 7.6± 2.2 | 77 (43.7)<br>0.13 | **132 (75.0)**<br>**0.03** |
| Aching<br>p | 78.5* | 7.0±2.1 | 86 (48.0)<br>0.37 | 130 (72.6)<br>0.47 |
| Gnawing<br>p | 35.1 | 7.3± 2.5 | 32 (40.0)<br>0.15 | 56 (70.0)<br>0.71 |
| Tender<br>p | 62.7 | 7.0±2.3 | **57 (39.8)**<br>**0.01** | 108 (75.5)<br>0.08 |
| Heavy<br>p | 45.2* | 6.8±2.2 | 47 (45.6)<br>0.81 | 72 (69.9)<br>0.63 |
| PARESTHESIA/<br>DYSTHESIA |  |  |  |  |
| Tingling<br>p | 23.2* | 5.9±2.4 | 24 (45.2)<br>0.84 | 37 (69.8)<br>0.76 |
| Numb<br>p | 19.7* | 6.0±2.7 | 17 (37.7)<br>0.19 | **38 (84.4)**<br>**0.03** |
| Pins and needles<br>p | 22.8* | 7.5± 2.1 | 25 (48.0)<br>0.79 | 33 (63.4)<br>0.14 |
| Cutting<br>p | 17.1 | 7.4± 2.2 | 20 (51.2)<br>0.51 | 29 (74.3)<br>0.66 |
| Pulling<br>p | 24.6 | 6.9± 2.2 | 22 (39.2)<br>0.21 | 41 (73.2)<br>0.74 |
| Cold/Freezing | 11.4 | 7.0± 2.4 | 13 (59.0)<br>0.13 | 16 (72.7)<br>0.94 |
| Hot/Burning<br>p | 24.3 | 7.5±3.5 | 19 (40.4)<br>0.57 | 41 (87.2)<br>0.01 |
| EVOKED |  |  |  |  |
| Biting/Chewing<br>p | 88.6* | 8.3± 1.9 | 96 (47.5)<br>0.38 | 152 (75.2)<br>0.00 |
| Eating/Drinking<br>Cold<br>p | 68.4* | 7.9±1.9 | 87 (55.7)<br>0.000001 | 113 (72.4)<br>0.64 |
| Eating/Drinking<br>Hot<br>p | 43.9 | 7.4± 2.2 | 52 (52.0)<br>0.14 | 74 (74.0)<br>0.46 |
| OTHER |  |  |  |  |
| Sharp<br>p | 49.6* | 8.0± 1.8 | 57 (50.4)<br>0.24 | 82 (72.5)<br>0.72 |
| Dull<br>p | 43.3* | 5.8± 2.3 | 47 (46.5)<br>0.99 | 67 (66.3)<br>0.12 |

n (%): number of patients (percentage of patients within the population N= 228) with the clinical finding that also reported the pain descriptor
\*Pain descriptors selected for analysis by 2-way ANOVA
\*\*Mean and standard deviations were calculated with patients who reported pain more than 0 on a NRS from 0-10.

We next wanted to determine if there was any independent association between reporting of a pain descriptor and two key sensory tests performed for endodontic diagnosis: a cold detection test (does the subject feel cold applied to the tooth), and a percussion sensitivity test (is tapping the tooth painful). The univariate analysis showed that patients describing pain as shooting, bursting and tender as well as pain evoked while eating or drinking cold things were more likely to respond positively to cold detection test, suggesting vital inflamed dental pulp mediates these sensations. Also, patients who described their pain as throbbing, numb, hot/burning and pain evoked by biting and chewing were more likely to experience pain upon mechanical percussion of the tooth (p < 0.05) (Table 3).

Among these 22 pain descriptors, those which had little relevance to tooth pain based on infrequent reporting or caused confusion to subjects, i.e. poor face validity (gnawing, tender, cutting, pulling, cold/freezing, hot/burning) were dropped from further analysis. The remaining descriptors intensity measures as well as global pain scores were further analyzed by 2-way ANOVA to assess the association with the independent clinical variables cold detection and percussion hypersensitivity (Table 5). Before proceeding with 2-way ANOVA, it was confirmed that age, sex and ethnicity were equally distributed among groupings by cold detection +/- and percussion hypersensitivity +/- (Table 4).

**4.** Distribution of Patient Characteristics Among Groups

4. Distribution of Patient Characteristics Among Groups
|  | negative cold/normal percussion | negative cold/painful percussion | positive cold/normal percussion | positive cold/painful percussion | p value |
| --- | --- | --- | --- | --- | --- |
| Age | 44 | 43 | 37 | 40 | 0.83 |
| Female n (%) | 13 (56.5) | 51 (51.5) | 21 (50.0) | 36 (56.2) | 0.89 |
| Hispanic or Latino n (%) | 3 (13.0) | 23 (23.2) | 15 (35.7) | 12 (18.7) | 0.13 |
| Race n (%) |  |  |  |  |  |
| White/Caucasian | 11 (47.8) | 46 (46.4) | 18 (42.8) | 32 (50.0) | 0.38 |
| Black/African American | 6 (26.0) | 16 (16.1) | 5 (11.9) | 15 (23.4) |  |
| Other | 6 (26.0) | 37 (37.3) | 19 (45.2) | 17 (26.5) |  |

n (%), number of patients (percentage of patients within the population N= 228)

**5.**
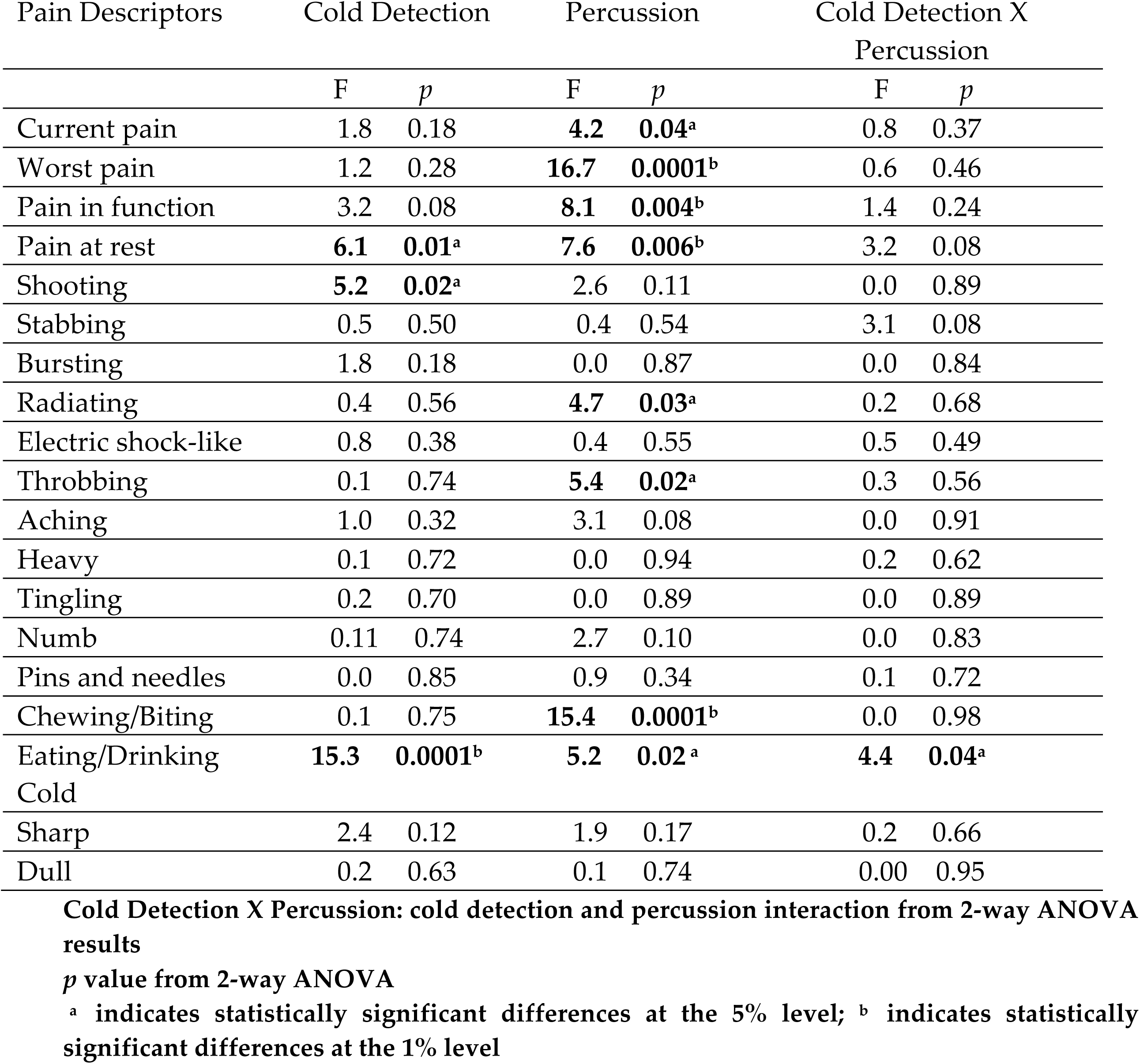
Pain Descriptors ANOVA Results

Patients with percussion hypersensitivity reported significantly higher levels of current pain, worst pain, pain at rest and pain in function. Patients with positive cold detection reported higher levels of pain at rest; i.e. spontaneous pain. There was a significant interaction between the cold responsiveness and percussion hypersensitivity on the intensity of pain evoked by cold. Patients with a positive cold detection and percussion hypersensitivity experienced more intense evoked pain with eating or drinking cold things. Patients with positive cold detection reported higher intensity shooting pain. Finally, patients with percussion hypersensitivity reported higher levels of radiating, throbbing pain and pain evoked by chewing/biting (Table 5) (Figure 2).

**Figure 2.**
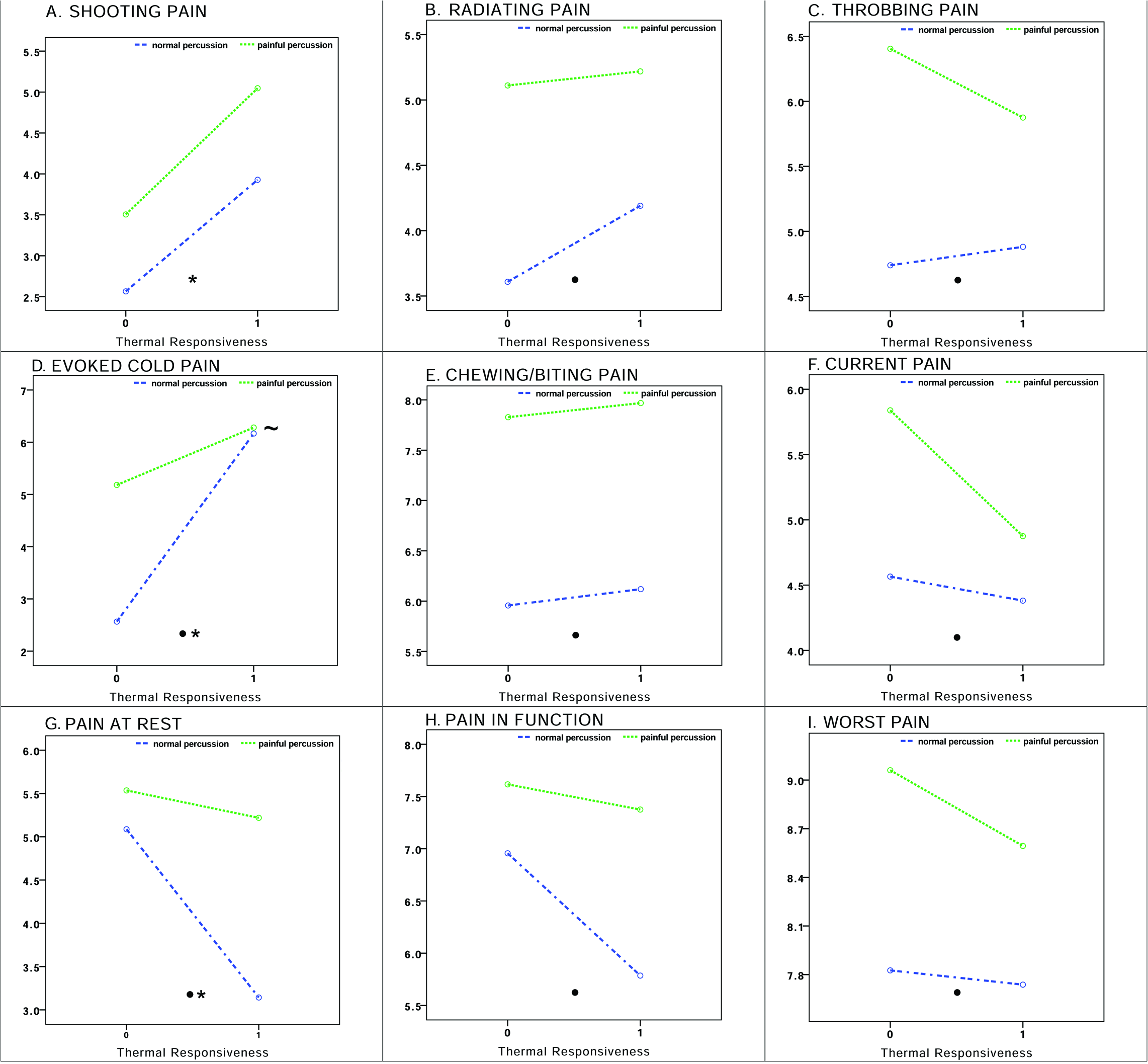
A: Patients with positive cold detection reported higher intensity shooting pain. B, C, E: Patients with percussion hypersensitivity reported higher levels of radiating, throbbing pain and pain evoked by chewing/biting. D: There is a significant interaction between the cold responsiveness and percussion hypersensitivity on the intensity of pain evoked by cold. Patients with positive cold detection and percussion hypersensitivity experienced more intense evoked pain with eating or drinking cold things. F, G, H, I: Patients with percussion hypersensitivity reported significantly higher levels of current pain, worst pain, pain at rest and pain in function.

## Discussion

Preoperative pain assessment is important for understanding the burden of tooth pain in patients (Nusstein and Beck 2003; Toure et al. 2007) as well as for the identification of pulpal and periapical pathology (Levin et al. 2009). What is more, preoperative pain is the most important predictor of post-operative pain and even chronic pain after treatment (Kehlet et al. 2006; Law et al. 2015; Polycarpou et al. 2005). Because obtaining a detailed preoperative pain history is the crucial part of our study, we focused on emergency patients like in the studies where preoperative pain is evaluated (Oguntebi et al. 1992; Toure et al. 2007). In this study, we report the intensity and quality of preoperative pain patients with toothache and their relation with clinical findings. Our results suggest that pain descriptors indeed associate with the clinical status of pulpal and periapical tissues.

In this study, patients who visited the emergency dental clinic with tooth pain had average pain intensity around 5 on a 10-point numerical rating scale (NRS), suggesting a moderate level of pain. The pain intensity reported was higher when describing pain in function (7.5 on NRS), and even higher when describing worst pain (8.5 on NRS). These pain levels are severe, and highlight the extreme burden of dental pain for some patients. These results are similar to previous reports in the literature. (Rechenberg et al. 2016; Toure et al. 2007) As appropriate dental care, including endodontic treatments such as root canal or pulpotomy, or extraction in non-restorable teeth, are highly effective treatments for quickly relieving this pain (Law et al. 2014), these findings highlight that it is essential for everyone has access to affordable dental care.

This study presents data that not only on intensity of the preoperative pain but also on the duration of it causing a burden on the study population. It is found out that cumulatively 59% of the patients had pain less than a week which is similar to the study by Rechenberg; where it was also found out that 45.7% of the patients diagnosed with irreversible pulpitis, 69.9% of patients diagnosed with acute apical periodontitis waited less than a week before attending to dental emergency (Rechenberg et al. 2016). However, we also found out that about one fifth of patients (22%) had pain for one month or more which is significant because literature states that the longer the period of time with preoperative pain the more likely to develop chronic pain after endodontic treatment (Polycarpou et al. 2005).

In terms of pain medication use, it is found out that for pain relief, 72% of the patients had used some type of medication, with 16% taking opioids. These findings are similar to those of Nusstein and Beck, where 81–83% of presenting symptomatic patients had taken analgesics with and 20–23% choosing opioid analgesics (Nusstein and Beck 2003). This is important given that opioids are much less effective than non steroidal anti-inflammatory drugs (NSAIDs), and many/most of these patients are using the analgesics from friends or family members prescribed them for another purpose (Aminoshariae et al. 2016). Of course, given the magnitude and terrible impact of the current opioid crisis in the US, it is of course important to address how many people are getting exposed to opioids due to attempts to mitigate untreated dental pain. (Table 2).

We found that subjects who report pain upon percussion of the affected tooth reported higher levels of pain at rest, pain at function, present pain and worst pain (p < 0.05) (Table 5). This is consistent with other work demonstrating that toothache patients without percussion hypersensitivity waited longer than patients diagnosed with percussion hypersensitivity before going for dental treatment (Toure et al. 2007) Another study showed, 92% of the patients who attended emergency clinic with pain had percussion hypersensitivity regardless of their diagnosis which is comparable to and even higher than our finding of 71.5% (Rechenberg et al. 2016). Together this supports that mechanical hypersensitivity of the periapical tissues strongly impacts the pain experience of toothache and, as such, is a major factor in motivating dental visits for treatment.

Traditionally, percussion hypersensitivity was thought to identify that infection or inflammation has spread beyond the confines of the tooth to the periapical tissues (Ingle JI 2008). More recent studies point out percussion hypersensitivity may identify mechanical allodynia due to central sensitization (Latremoliere and Woolf 2009; Owatz et al. 2007; Pigg et al. 2016). Models of pulpitis, in which the inflammation in confined to the pulp, causes changes in the nucleus caudalis including upregulation of p38 MAPK, increased expression of microglia and astrocytes and central sensitization (Worsley et al. 2014; Xie et al. 2007; Zhang et al. 2006). These changes in the central nervous system can contribute to mechanical hypersensitivity, which manifests as painful percussion on a clinical exam. Therefore, we and others speculate that the reason why patients with percussion sensitivity experienced preoperative pain more intensely could be explained by lowered mechanical thresholds due to central sensitization (List et al. 2008; Nikolajsen et al. 2000; Nikolajsen and Jensen 2001; Owatz et al. 2007). Of course in some cases, in more advanced infections, the inflammation has spread to the periapical tissues and inflammatory mediators could directly cause hypersensitivity of periapical afferents, via peripheral sensitization.

Pain descriptors can help clinicians in diagnosing certain categories of pain such as neuropathic pain, and in some cases might reveal the underlying biological mechanisms of the disease (Bouhassira et al. 2005). We found that dental pain patients with hypersensitivity to percussion reported higher levels of radiating and throbbing pain (p<0.05) (Table 5). Throbbing pain is specified deep as opposed to superficial pain (Victor et al. 2008). In two consecutive case studies, the underlying biology of throbbing pain in migraine is questioned. It was found out that throbbing pain is not correlated with arterial pulse (Ahn 2010) but was synchronized with alpha waves measured with EEG pointing out throbbing pain may tell us about the central processing of pain (Mo et al. 2013).

Similarly, patients with percussion hypersensitivity reported higher levels of pain when chewing and biting. This is an expected result since percussion, biting and pressure all test for mechanical allodynia (Khan et al. 2007). What is more, a significant interaction between the effects of cold detection and percussion hypersensitivity on the intensity of evoked pain by cold was found (p<0.05) (Table 5). This finding is actually in line with the studies where percussion hypersensitivity is present with patients diagnosed with irreversible pulpitis (positive cold detection) (Owatz et al. 2007; Rechenberg et al. 2016) where it is hypothesized that this could be in fact due to central sensitization (Khan et al. 2007; Owatz et al. 2007).

Patients with a positive response to cold detection more frequently used the pain descriptors “shooting” and “bursting” and also reported higher intensity shooting pain. These pain descriptors describe paroxysmal pain, and this finding is consistent with other reports that describe pulpal inflammation as producing paroxysmal type pain (Yair Sharav 2010). The dental pulp is a uniquely innervated tissue with the majority of fibers being medium to large size myelinated fibers, and lacking certain classes of afferents including IB4 expressing c-fibers (Fried et al. 2011; Gibbs et al. 2011). Perhaps the unique neurobiological signature of pulp innervating afferents causes the unique sensory experience of pulpitis, which includes very intense paroxysmal type pain.

Another interesting finding of this study is that dental pain patients chose pain descriptors that are associated with neuropathic pain more frequently than anticipated. Although not associated with a specific clinical test, 23% of study participants chose the descriptors “pins and needles” and “tingling” to describe their tooth pain. Further, patients with percussion pain; i.e. mechanical allodynia of the affected tooth, more frequently reported their pain felt “numb” and “burning” (Bouhassira et al. 2005; Lopez-Jornet et al. 2017). All of these descriptors are commonly associated with neuropathic pain, in which there is some disease or damage to the sensory nervous system itself. As pulpitis progresses, die back of pulp innervating afferents occurs as the pulp tissue succumbs to infection. The association of reporting neuropathic pain descriptors and observing pain on percussion suggests that there could be a neuropathic component of painful toothache. This is also consistent with animal studies which have found markers of nerve injury upregulated after pulp injury. As 3–10% of patients experience persistent pain after dental interventions such as root canals, these findings raise the question whether the neuropathy might occur before the dental intervention even takes place (Nixdorf et al. 2016; Polycarpou et al. 2005; Vena et al. 2014). Also, do patients experiencing pre-operative neuropathic type pain have prolonged post-operative or even persistent pain after root canal treatment. Further studies are needed to answer these important questions.

The first strength of this study is that it reveals detailed preoperative pain data which emphasizes how preoperative dental pain is needed to be taken into account meticulously. The second strength of the study is that patient reported pain descriptors may differentiate different stages of pulpal and periapical pain.

There are also limitations in this study. The first limitation is that tooth vitality is determined by surrogate tests instead of a more objective method of inspection of blood when the pulp chamber was exposed. In this study in order to access to more patients, and to obtain more generalizing results, patients seen by endodontic residents were included and this made it difficult to obtain data that needed to be controlled more meticulously like observing bleeding when the pulp chamber was exposed.

The major limitation in this study is that pain data collected from patients is subjective. What is more, although thermal and mechanical tests are thought to be more objective; their results are also obtained by asking the patients which again results in a subjective data. However, we still don’t have biomarkers in endodontic practice that we can rely on as objective findings to distinguish the biological state of the pulp and periapical tissues (Rechenberg 2014). Therefore, today, in endodontic practice, we need to obtain detailed subjective data and interpret them together to understand the underlying biology of pulpal and periapical pain.

